## supplementary data for "Structure of human DPPA3 bound to the UHRF1 PHD finger reveals its functional and structural differences from mouse DPPA3"

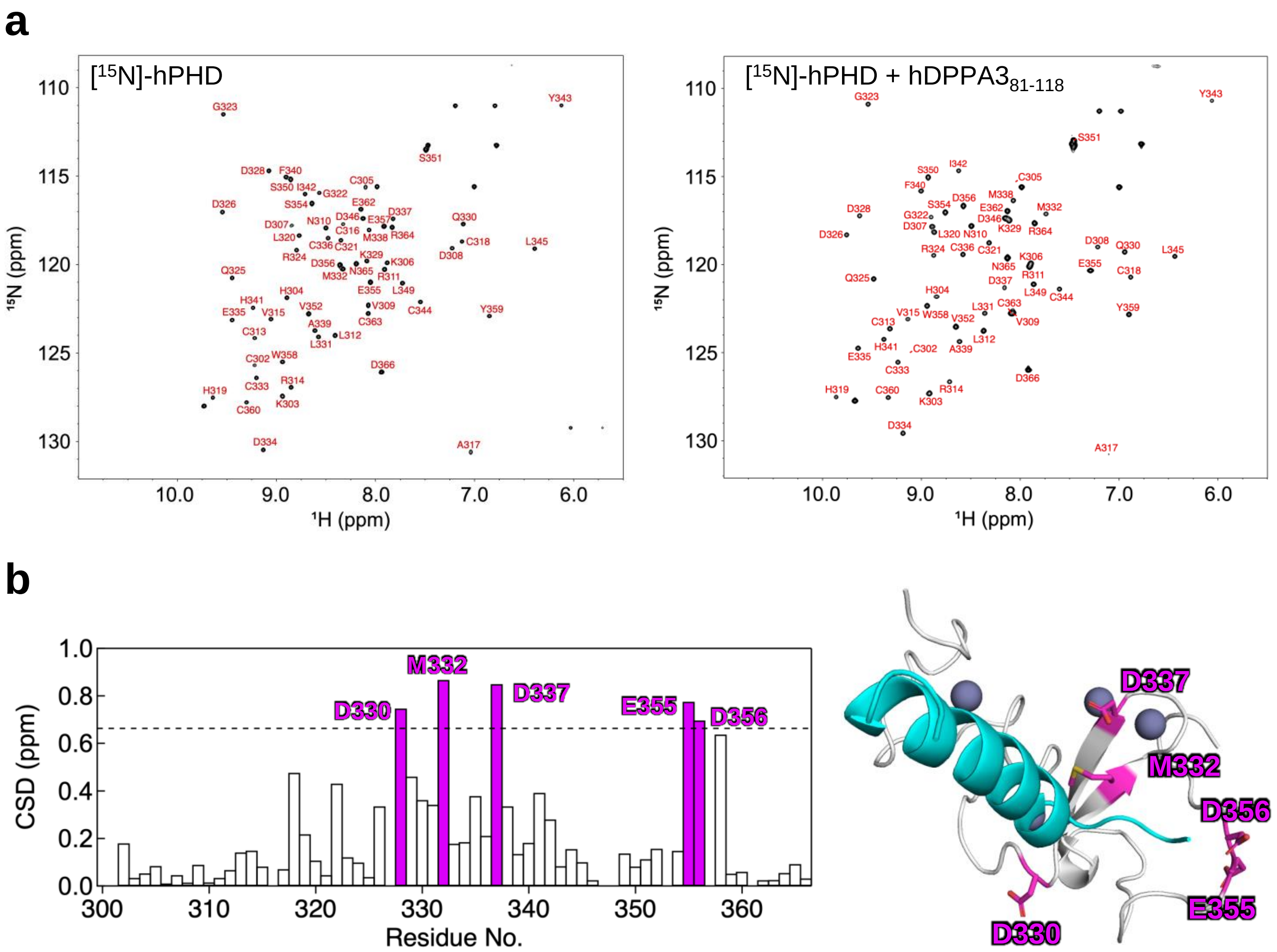


**Supplementary Figure 1: Nuclear magnetic resonance (NMR) titration experiments**

**(a)** Signal assignments in the $^{1}H$-$^{15}N$ HSQC spectra of hPHD in the free state (left) and complex state with hDPPA3$_{81-118}$ (right). **(b)** Weighted average chemical shift differences in $^{1}H$ and $^{15}N$ resonances between free hPHD and hPHD in complex with hDPPA3$_{81-118}$ (left). The dashed line represents the mean plus 2 standard deviations. Mapping of D330, M332, D337, E355, and D356 in the crystal structure (right).

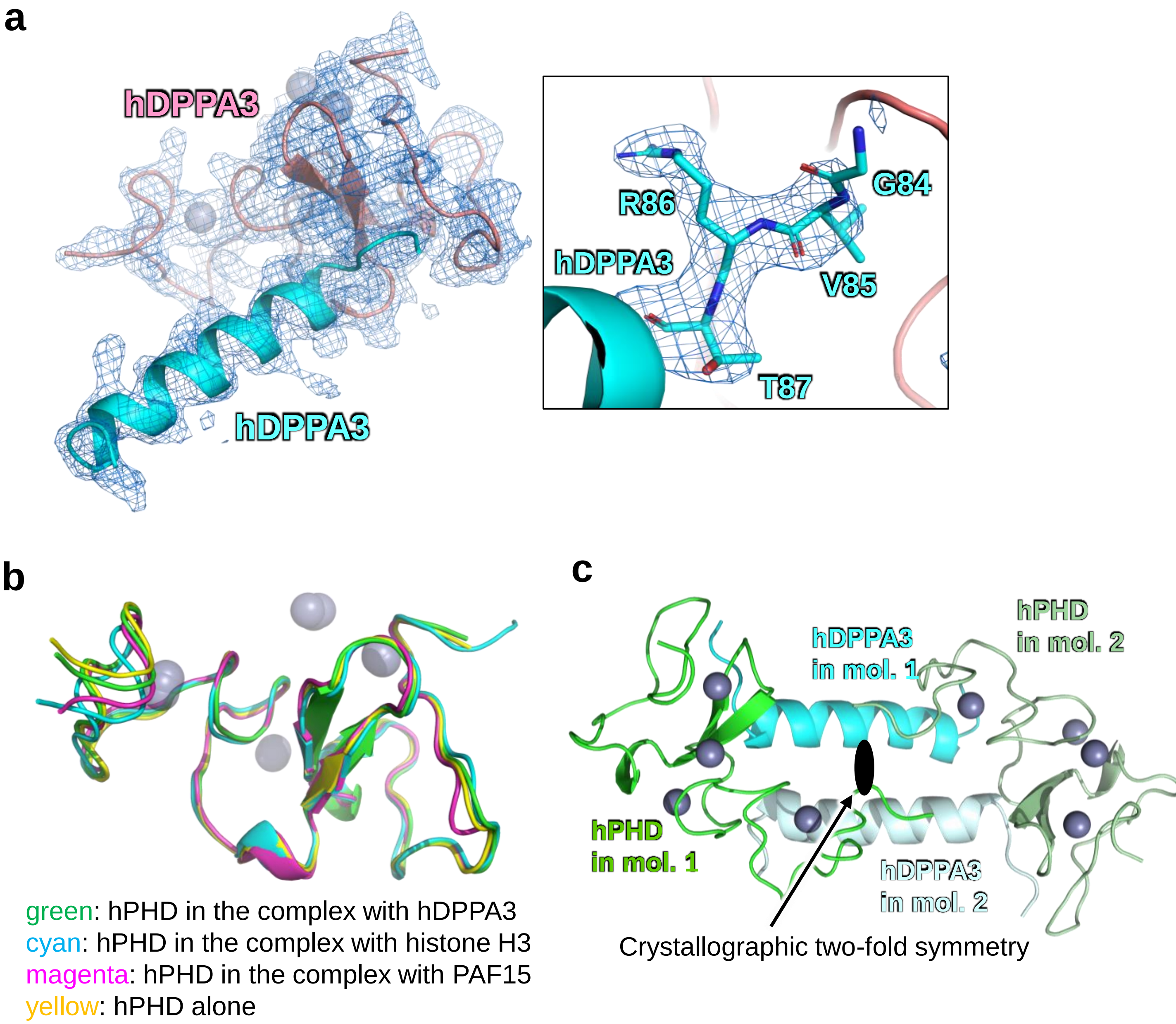


**Supplementary Figure 2: Structural features of the hPHD:hDPPA3 complex in crystal form**

**(a)** Overlay of 2|*F*o|-|*F*c| map countered at 1.0 σ (blue mesh) on the cartoon model of the hPHD:hDPPA3$_{81\text{-}118}$ complex. The figure on the right shows a close-up view of the VRT motif of hDPPA3. The Omit map contoured at 3 σ is shown as a blue mesh. **(b)** Structural comparison of the human UHRF1 PHD finger in the complex with hDPPA3 (green), histone H3 (cyan), and PAF15 (magenta) and of the structure of hPHD alone (yellow). **(c)** The hPHD:hDPPA3$_{81\text{-}118}$ complex forms a dimer with a crystallographic symmetry-related molecule via interaction with the α-helix in hDPPA3. The crystallographic two-fold symmetry symbol is indicated.

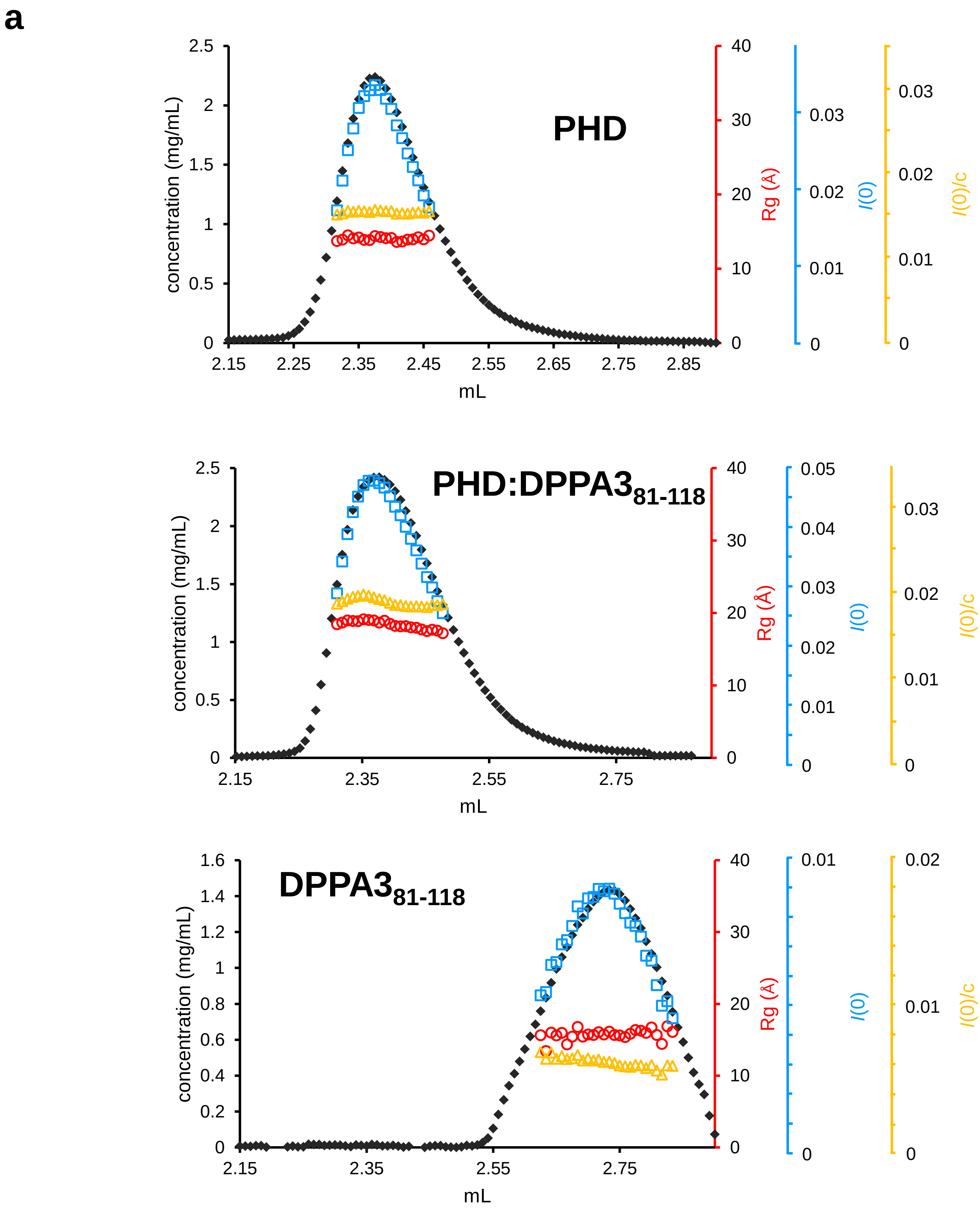
a
PHD
concentration (mg/mL)
Rg (Å)
I(0)
I(0)/c
mL
PHD:DPPA3$_{81-118}$
DPPA3$_{81-118}$

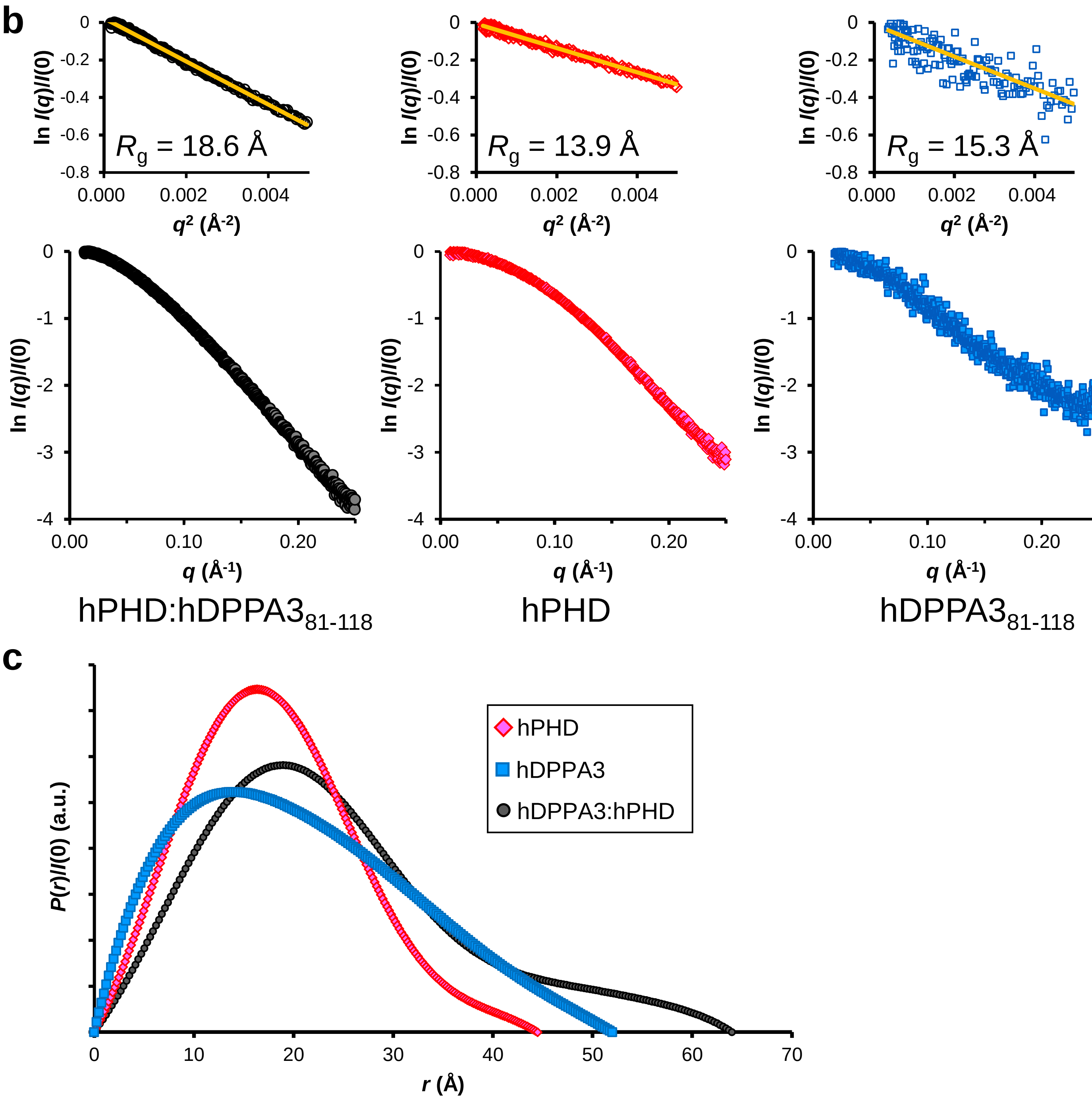


**Supplementary Figure 3: Size-exclusion chromatography–small-angle X-ray scattering (SEC-SAXS) data.**

**(a)** SEC-SAXS data of hPHD:hDPPA3$_{81\text{-}118}$ complex (top), hPHD (middle), hDPPA3$_{81\text{-}118}$ (bottom). Absorption at 280 nm, $I(0)$ calculated from the Guinier analysis, $R_g$ values, and $I(0)/c$ (concentration: mg/ml) are shown as black diamonds, cyan squares, red circles, and yellow triangles, respectively. **(b)** SAXS scattering intensity of the hPHD:hDPPA3$_{81\text{-}118}$ complex (left), hPHD (center), and hDPPA3$_{81\text{-}118}$ (right). The top panels indicate the Guinier plot with $qR_g < 1.3$ shown as an orange line. $R_g$ values are shown in the graphs. **(c)** Overlay of the distance distribution function, P(r), of hPHD:hDPPA3$_{81\text{-}118}$ complex (black), hPHD (pink), and hDPPA3$_{81\text{-}118}$ (cyan).

a

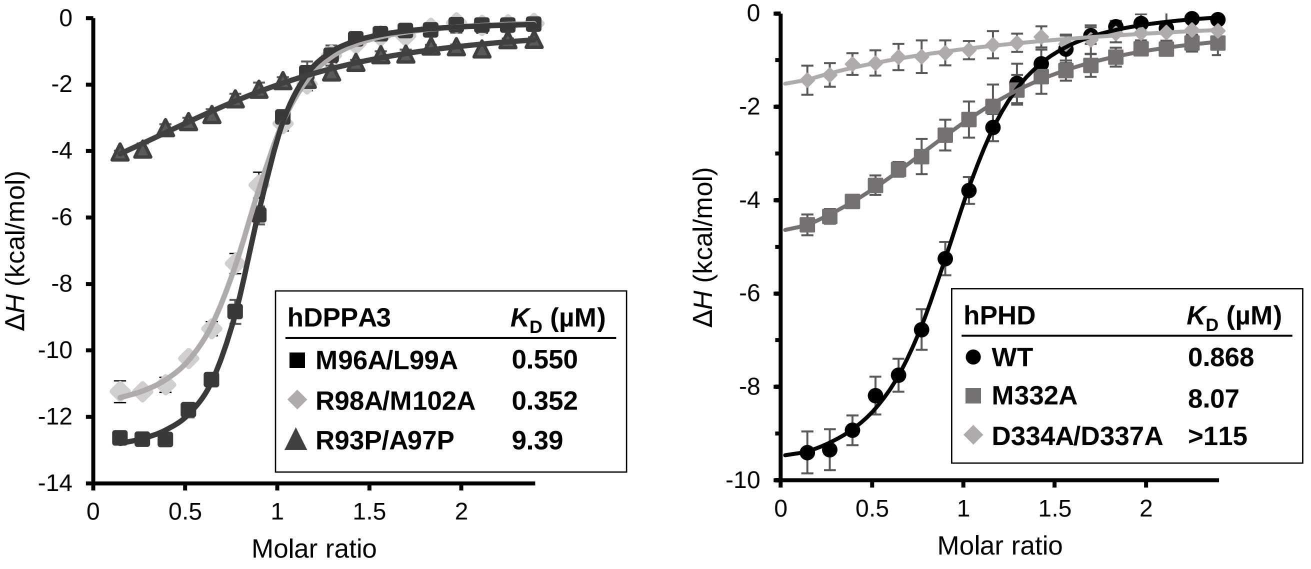


b

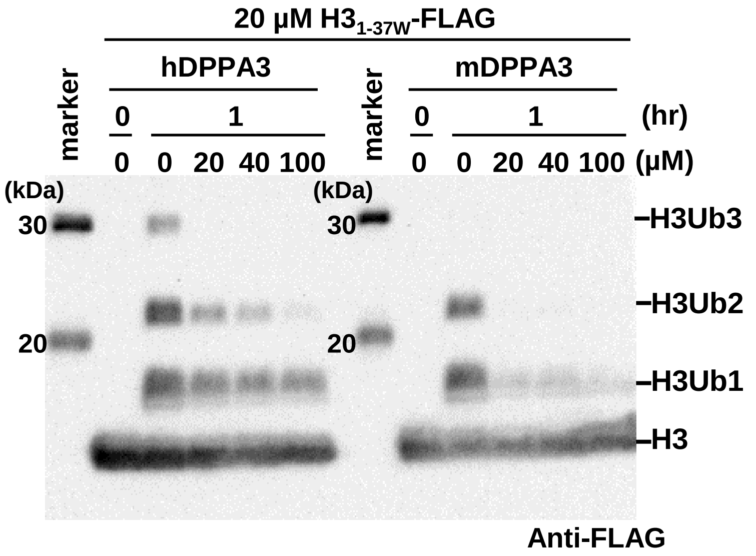


**Supplementary Figure 4: Biochemical assays.**

**(a)** Isothermal titration calorimetry (ITC) measurements for mutants of $hDPPA3_{81\text{-}118}$ and hPHD. Superimposition of enthalpy change plots with standard deviations. Data are presented as mean ± SD for n = 3. **(b)** An in vitro ubiquitination assay to compare the inhibitory effects of hDPPA3 and mDPPA3. The gel image is representative of n = 3 independent experiments.

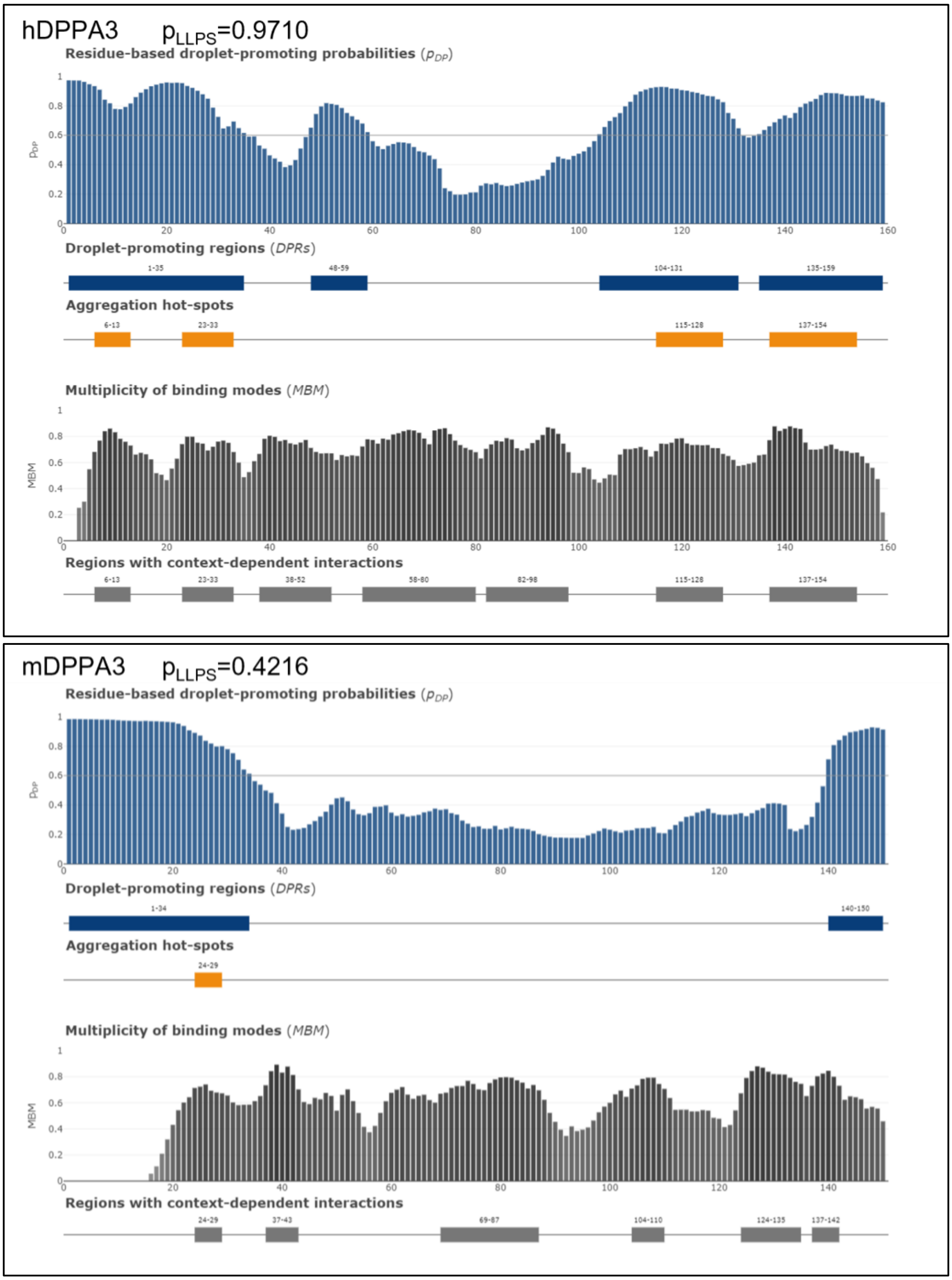


**Supplementary Figure 5: Prediction of droplet formation in human (upper panel) and mouse DPPA3 (lower pannel).**

PLLPS is the probability of forming a droplet state through liquid-liquid phase separation. PLLPS ≥ 0.60 are assigned as droplet-drivers.

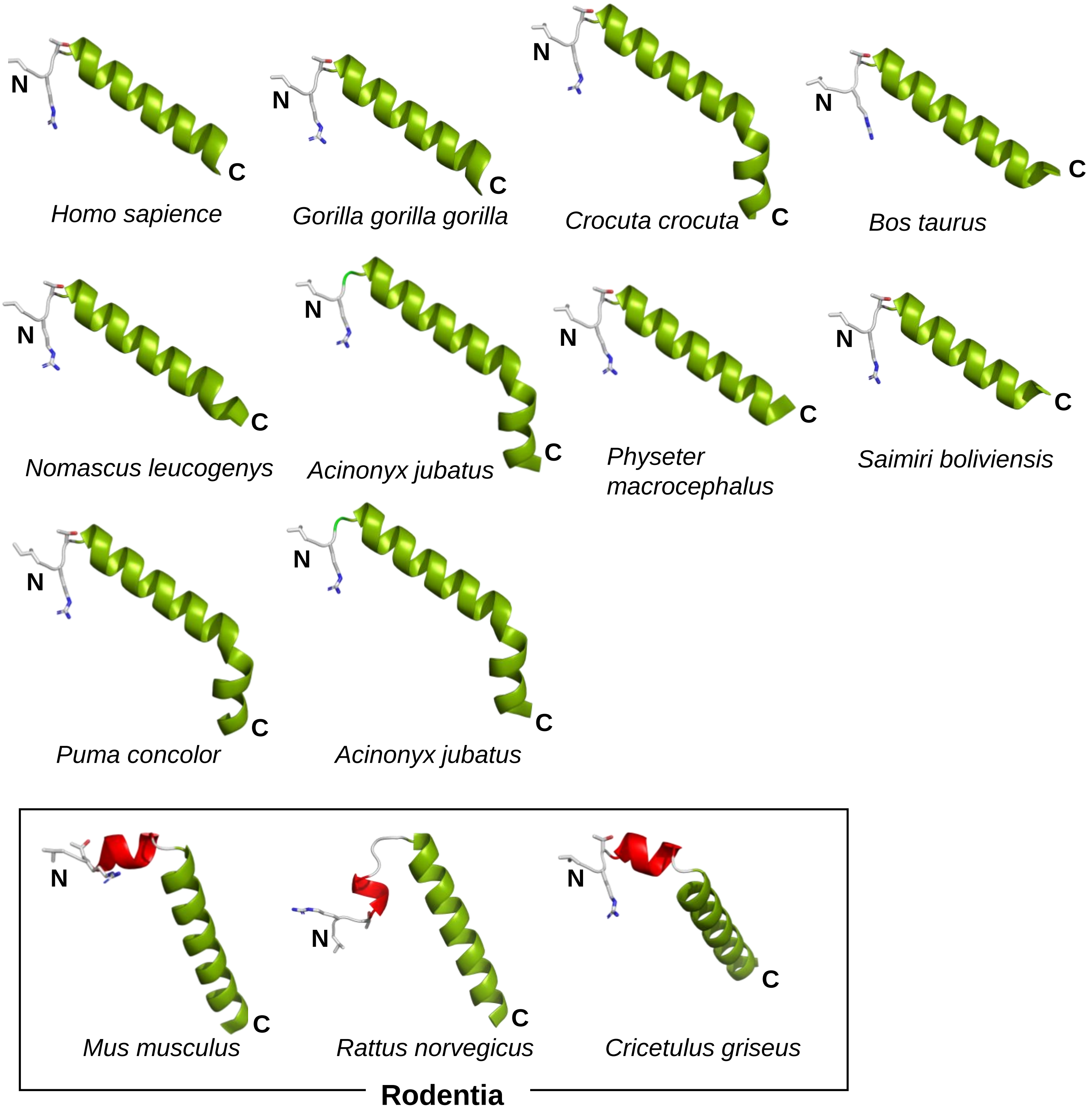


**Supplementary Figure 6: AlphaFold 2 structural prediction of DPPA3 in various species.**
The VRT motif and the following α-helix of DPPA3, predicted by AlphaFold2, are displayed. White sticks indicate side chains of the conserved VRT motif. α-Helices are shown as cartoon models, in which the long α-helix is colored green and the short α-helix, unique to Rodentia, is displayed in red.

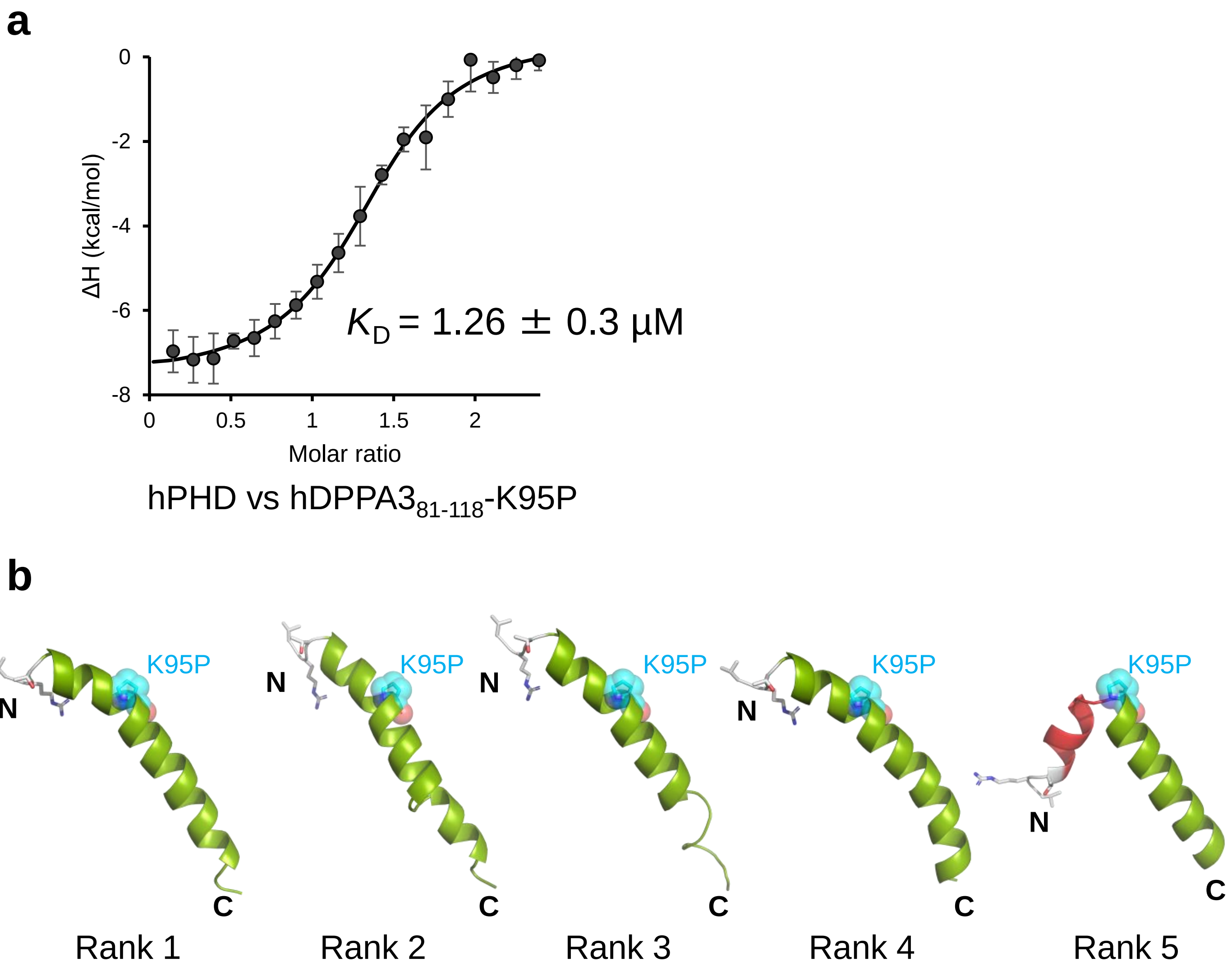


**Supplementary Figure 7: Analysis of K95P mutant of hDPPA3.**
**(a)** ITC measurements of the K95P mutant hDPPA3$_{81-118}$ and WT hPHD. Superimposition of enthalpy change plots with standard deviations. Data are presented as mean ± SD for n = 3. **(b)** AlphaFold2 structure prediction of K95P mutant of hDPPA3$_{81-118}$. K95P is shown as a cyan stick superimposed on a transparent sphere. The VRT motif is depicted as a stick model.

**Supplementary Table 1**

**Sample details**

| Sample name | **hPHD** | **hPHD:hDPPA3** | **hDPPA3** |
|---|---|---|---|
| Organism | *Homo sapiens sapiens* | | |
| UniProt sequence ID | Q96T88 (UHRF1) | Q96T88 (UHRF1)<br>Q6W0C5 (DPPA3) | Q6W0C5 (DPPA3) |
| Extinction coefficient ε | 8480 | 9970 | 1490 |
| Calculated monomeric Mr from sequence (kDa | 7.8 ($PHD_{299-366}$) | 12.2:<br>[7.8 ($PHD_{299-366}$) + 4.4<br>($DPPA3_{81-117}$)] | 4.4 ($DPPA3_{81-117}$) |
| HPLC system | Nexera/Prominence-I (Shimazu) | | |
| SEC Column | Superdex™ 200 Increase 5/150 GL | | |
| Temperature (K) | 293 | | |
| Injection volume (µL) | 50 | | |
| Loading concentration (mg/mL) | 12 | | |
| Flow rate (mL/min) | 0.025 | | |
| SEC buffer | 20 mM Tris-HCl (pH 7.5), 150 mM NaCl, 2 mM DTT, 10 µM Zinc acetate and 5% glycerol | | |

**Data-collection parameters**

| | | | |
|---|---|---|---|
| Beamline | Photon Factory BL-10C | | |
| Detector | Pilatus3 2 M | | |
| Sample-to-detector distance (mm) | 2,082 | | |
| Wavelength (Å) | 1.5 | | |
| $q$ range ($Å^{-1}$) | 0.0083 - 0.264 | 0.00865 - 0.264 | 0.018-0.264 |
| Exposure time (s) | 20/frame | | |
| Flux (photons/s) | $1.1\times10^{11}$ | | |
| Beam size | 0.63 mm (H)×0.18 mm (V) | | |
| Concentration range (mg/mL) | 1.17 - 2.18 | 1.3 - 2.46 | 0.739-1.44 |
| Absolute scaling method | Using the scattering intensity of water | | |
| Normalization | To transmitted intensity by beam-stop counter | | |

**Structural parameters**

| | | | |
|---|---|---|---|
| **Guinier analysis** | | | |
| $I(0)$ ($cm^{-1}$) | $0.015 \pm 1.8\ E^{-5}$ | $0.019 \pm 2.2\ E^{-5}$ | $0.0057 \pm 9.0\ E^{-5}$ |
| $R_g$ (Å) | 13.9 ± 0.03 Å | 18.6 ± 0.04 Å | 15.3 ± 0.43 Å |
| $q$-range ($Å^{-1}$), point range | 0.18-1.30, 14-248 | 0.20-1.30, 7-179 | 0.30-1.30, 3-186 |
| ***P*(r) analysis** | | | |
| $I(0)$ ($cm^{-1}$) | $0.015 \pm 8.6\ E^{-5}$ | $0.019 \pm 1.2\ E^{-4}$ | $0.0058 \pm 1.1\ E^{-4}$ |
| $R_g$ (Å) | 14.0 ± 0.11 Å | 19.1 ± 0.19 Å | 16.5 ± 0.32 Å |
| $D_{max}$ (Å) | 44.5 | 64.0 | 52.0 |
| $q$-range ($Å^{-1}$), point range | 0.0128-0.2642, 14-742 | 0.0100-0.2156, 5-600 | 0.0211-0.2642, 10-714 |
| Porod volume estimate ($Å^3$) | 16393 | 16873 | 13315 |
| Dry volume calculated from sequence ($Å^3$) | 9381 | 14711 | 5352 |
| Partical specific volume ($cm^3g^{-1}$) | 0.743 | 0.743 | 0.743 |
| Molecular mass Mr [from $V_c$] (kDa) | 9.0 | 13.0 | 7.9 |
| Calculated monomeric Mr from sequence (kDa | 7.8 | 12.2 | 4.4 |

**Software employed**

| | |
|---|---|
| Primary data reduction | Sangler 2.1.39 |
| Guinier Analysis | AutoGuinier (ATSAS 3.1.3) |
| Zero-concentration Extrapolation | MOLASS 1.0.10 |
| Data processing | PRIMUS |
